# Striatal cholinergic receptor activation causes a rapid, selective, & state-dependent rise in corticostriatal *β* activity

**DOI:** 10.1101/148551

**Authors:** Benjamin R. Pittman-Polletta, Allison Quach, Ali I. Mohammed, Michael Romano, Krishnakanth Kondabolou, Nancy J. Kopell, Xue Han, Michelle M. McCarthy

**Affiliations:** Boston University, Department of Mathematics & Statistics, Boston, MA 02215; Boston University, Department of Biomedical Engineering, Boston, MA 02215

**Keywords:** oscillations, basal ganglia, motor disorders, phase-locking, acetylcholine

## Abstract

Cortico-basal ganglia-thalamic (CBT) *β* oscillations (15–30 Hz) are elevated in Parkinson’s disease and correlated with movement disability. To date, no experimental paradigm outside of loss of dopamine has been able to specifically elevate *β* oscillations in the CBT loop. Here, we show that activation of striatal cholinergic receptors selectively increased *β* oscillations in mouse striatum and motor cortex. In individuals showing simultaneous *β* increases in both striatum and M1, *β* partial directed coherence (PDC) increased from striatum to M1 (but not in the reverse direction). In individuals that did not show simultaneous *β* increases, *β* PDC increased from M1 to striatum (but not in the reverse direction), and M1 was characterized by persistent *β*-HFO phase-amplitude coupling. Finally, the direction of *β* PDC distinguished between *β* subbands. This suggests: (1) striatal cholinergic tone exerts state-dependent and frequency-selective control over CBT *β* power and coordination; (2) ongoing rhythmic dynamics can determine whether elevated *β* oscillations are expressed in striatum and M1; (3) altered striatal cholinergic tone differentially modulates distinct *β* subbands.

## INTRODUCTION

The striatal cholinergic system is emerging as a key mediator of parkinsonian motor symptoms. Chronic loss of dopamine diminishes M4 autoreceptor functioning on striatal cholinergic interneurons (sChIs), which normally function to regulate release of acetylcholine (ACh) [Ding et al. (2006)]. Increased levels of ACh occur in the striatum of both the 6-OHDA rat model of Parkinson’s disease (PD) as well as after acute antagonism of dopamine D2 receptors [Ikarashi et al. (1997)]. Optogenetic activation of striatal cholinergic interneurons (SChIs) promotes parkinsonian-like motor symptoms in normal mice [Kondabolu et al. (2016)], whereas silencing SChIs improves motor symptoms in dopamine depleted parkinsonian mouse models [Maurice et al. (2015)]. Furthermore, intra-striatal infusion of cholinergic muscarinic antagonists alleviates parkinsonian motor symptoms [Ztaou et al. (2016)], and antipsychotic-induced parkinsonism has been recently attributed to SChI activity [Kharkwal et al. (2016)]. Despite the evidence linking the striatal cholinergic system and parkinsonian motor deficits, the neuronal dynamics mediating cholinergic-induced movement disability are unknown. Understanding the neuronal and network mechanisms underlying cholinergic-induced movement disability would provide a framework for identifying likely targets for therapeutic interventions.

A common measure of neuronal dynamics, the local field potential (LFP), shows elevated and coordinated oscillations within the extended *β* frequency range (8–35 Hz) in cortico-basal ganglia-thalamic (CBT) circuits in PD patients and animal models of PD that correlate with movement disability [Stein and Bar-Gad (2013); Kuhn et al. (2009, 2008, 2006); Mallet et al. (2008b); Sharott et al. (2005); Mallet et al. (2008a)]. To date, no experimental paradigm (pharmacologic or optogenetic) outside of loss of dopamine has been able to elevate oscillations in the CBT loop specifically in the *β* frequency range. While there is some suggestion that the striatal cholinergic system can modulate *β* oscillations in CBT circuits, evidence is lacking for specificity of modulation to the *β* frequency range. Previous studies optogenetically stimulating striatal cholinergic interneurons in normal mice elevated oscillatory power at all frequencies greater than approximately 10 Hz, including both *β* and broadband *γ* oscillations [Kondabolu et al. (2016)]. Prior studies with carbachol-induced *β* oscillations in striatum did not address specificity of elevation to the *β* frequency range [McCarthy et al. (2011)]. Additionally, since the recordings were limited to striatum, these studies could not determine more widespread effects of striatal cholinergic modulation within CBT circuits [McCarthy et al. (2011)]. To determine whether modulations of striatal cholinergic tone are sufficient to elevate and coordinate oscillations specific to the *β* frequency range in normal, non-parkinsonian CBT circuits, we elevated striatal cholinergic tone with infusions of carbachol while making novel simultaneous recordings of LFPs from both striatum and M1. We found that acute increases in striatal cholinergic tone can both selectively increase *β* oscillations in striatum and M1 as well as increase coordination between these two regions specifically in the *β* frequency range when beta power is elevated in both regions. We additionally found two distinct CBT dynamic states: one which promotes increases in cortico-striatal beta power and coordination, and one that prevents beta elevation and coordination. Interestingly, the states are distinguished by differences in pre-infusion phase-amplitude coupling (PAC), which persist after carbachol infusion. Since these changes in oscillatory activity are acute, these findings have the important implication that the mechanism of exaggerated *β* oscillations in PD may stem from normal physiological processes in CBT circuits, rather than from plastic changes due to the chronic loss of dopamine. This paper presents evidence for the existence of cholinergically-modulated *β*-generating network mechanisms within the CBT circuits of normal mice as well as evidence that both the power and coordination of *β* oscillations can be regulated by striatal cholinergic receptors.

## METHODS

### Animal Preparations and Recordings

All animal procedures were approved by Boston University’s Institutional Animal Care and Use Committee (IACUC), and are in compliance with the National Institutes of Health Guide for the Care and Use of Laboratory Animals. 12 adult mice, C57B6 wild type (crossed between Emx1IRES-cre and Ai 35 mice, both obtained from Jackson Laboratories), or Emx-Arch transgenic mice (Jackson Laboratories), both males and females, were used in this study. Procedures were performed similarly to those previously described [McCarthy et al. (2011); Kondabolu et al. (2016)].

Stereotactic surgeries were performed under isoflurane general anesthesia. Analgesic buprenorphine was administered as postoperative care for at least 48 h. Mice were implanted with custom head-plates that allowed access to both striatum and motor cortex, and recording sites were marked. All recording and behavioral experiments were performed after the animals recovered from surgery, typically 3–6 d after the surgery. Upon recovery from surgery, mice were head-fixed and recorded from while awake, with glass electrodes (Warner instruments; G100F-4) filled with 0.9% saline solution (impedance 1-5 mOhm). Simultaneous recordings were made unilaterally in the striatum (stereotactic coordinate: AP (anterior posterior) = 0, ML (medial lateral) = 2.5-3 ML, and DV (dorsal ventral) = 2.0–2.5), and the primary motor cortex (M1, AP=-2, ML=1, DV=0.7-1). During recording, we also visualized glass electrode penetration depth and tracked neural activity patterns, and used them to verify electrode placement – in M1 below the pia, and in striatum beneath the corpus callosum, the latter being identified as an area with little neural activity. Local field potential was recorded with a multiclamp 700A amplifier (Molecular Devices), digitized with a Digidata 1440 (Molecular Devices, Inc.) at 20kHz, without additional filtering. Raw data were analyzed offline, and effectively high-pass filtered via downsampling (at 500 Hz) and low-pass filtered via detrending (at 0.05 Hz, see below).

The mice were habituated on the head fixed set-up for 2 days before infusions were performed. A custom cannula was coupled to the striatal recording electrode with the cannula tip positioned 500*μ*m above the electrode tip. Before carbachol infusion, baseline recordings were made for at least 5 minutes (7.9 ±2.3mins,meas.d., n = 12 mice). Carbachol (Sigma C4382) was dissolved in saline at a final concentration of 1mM, and infused at 0.2 *μ*L/min for 5mins (total 1 *μ*L). Recording was performed throughout the infusion period and for at least 12.5 minutes after infusion was started, with the average length of the recordings following the start of infusion being 24.6 ± 14.6 mins (mean ± s.d., n = 12 mice). Mice were not infused with carbachol in any recording sessions prior to those analyzed.

## Data Analysis

### Obtaining Band-Limited Power at Various Frequencies

Power increases observed after carbachol infusion were often bursty rather than continuous. To capture short-duration changes in spectral power, we calculated spectral measures using wavelets (Fig. 1a). Simultaneously recorded LFPs from each session were first detrended using a 20 second moving average, and then downsampled to 500 Hz sampling rate. They were then convolved with complex Morlet wavelets, spaced every 1 Hz between 1 and 200 Hz. The length of each wavelet was frequency dependent, with the time constant of exponential rolloff of the wavelet spaced linearly between 3 cycles at 1 Hz (for a resulting wavelet length of 3400 ms using eeglab function dftfilt3) and 21 cycles at 200 Hz (for a resulting wavelet length of 120 ms using eeglab function dftfilt3). The modulus of this complex signal provided the instantaneous amplitude or power for each frequency at each time point; its angle provided the instantaneous phase for each frequency at each time point. Power within each of 6 frequency bands – *δ* (1–4 Hz), *θ* (4–8 Hz), *α* /low *β* (8–15 Hz), *β* (15–30 Hz), *γ* (30–100 Hz), and high-frequency oscillation (HFO, 120–180 Hz) – were obtained as the square root of the summed squared power across each frequency range, at each time point.

**Figure 1:**
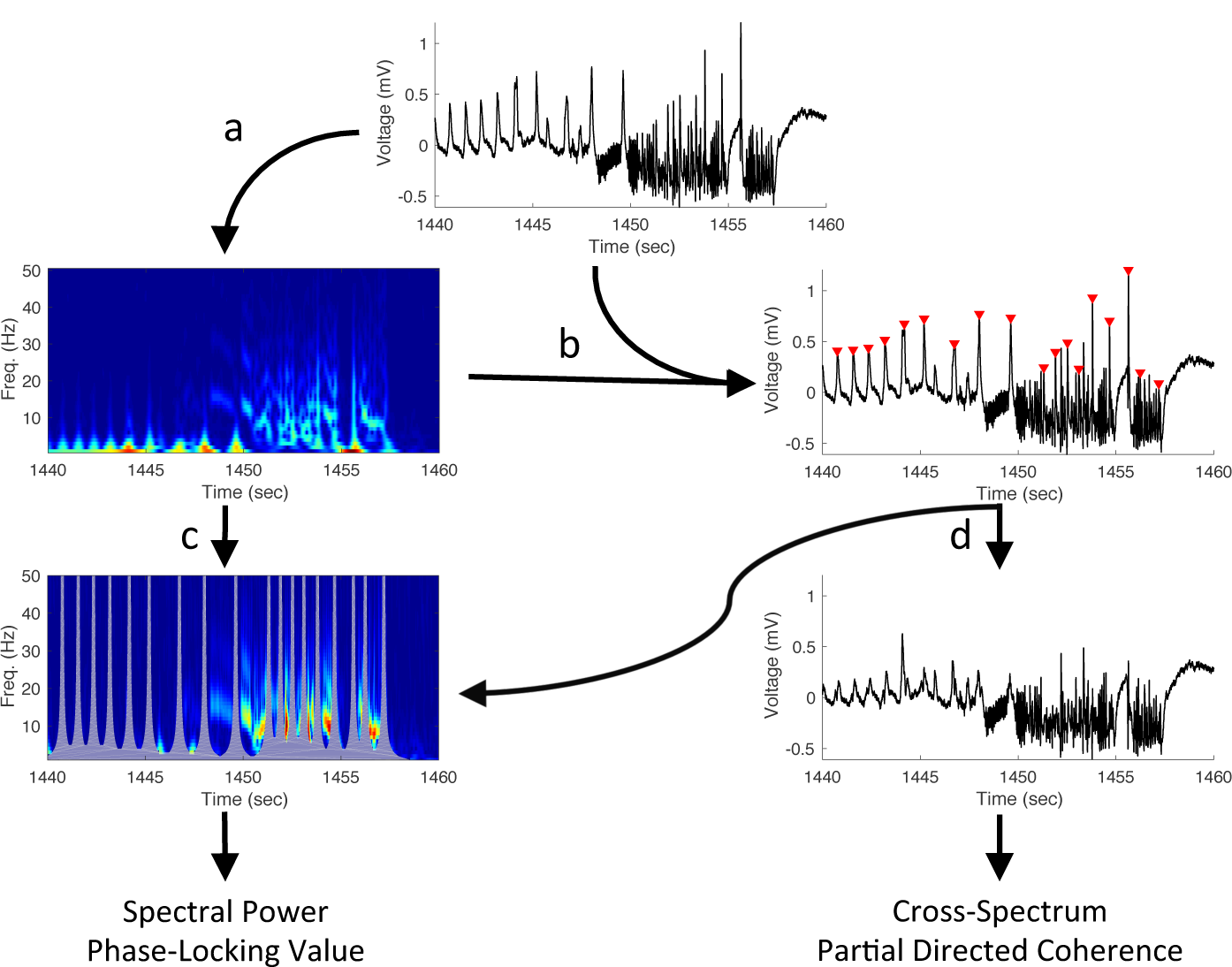
Peak artifacts were removed from spectral and time series data before analysis. Various stages of the peak-removal process are shown for a 20 second period in a representative post-carbachol recording. (a) Spectral information was extracted from LFP time series using complex Morlet wavelets. (b) Spectral and time series data were used to identify peak artifacts. (c) Spectral information potentially contaminated by peak artifacts was removed from measurements of spectral power and phase-locking value (indicated by gray overlays) for further analyses. (d) Peak artifacts were removed from time series data for analyses of cross-spectral power and partial directed coherence.

### Identifying Broadband LFP Deflections

Carbachol infusion often produced large peak-like LFP deflections (“peaks”, Fig. 1). These “peaks” had a broadband signature, with the most energy at lower frequencies, and thus complicated our analysis at specific frequency bands. To prevent the spectral signatures of these peaks from biasing our analyses, these “peaks” were first identified and removed before any spectral analysis.

Peak identification (Fig. 1b) was accomplished using one of two methods. For the first method, we first identified putative peak-like deflections using Matlab’s findpeaks function, which finds local maxima in the LFP traces that are separated by a minimum time, *t*_*min*_. This argument *t*_*min*_ (typically around 1 second) was determined by visual inspection for each recording to ensure that all possible artifactual peaks were selected. We then extracted the LFP waveform from *t*_*min*_ /2 before to *t*_*min*_ /2 after the identified putative peaks, and these waveforms were clustered using Matlab’s kmeans clustering function. The clusters corresponding to peaks were manually marked for removal. This method was successful in identifying peak-like deflections that exhibited highly stereotypical waveforms.

In the few cases when the first method failed due to peaks being insufficiently stereotypical, a second method was used. In this method, real peaks were selected from putative peaks by manually setting thresholds on “peak prominence”, a measure of how large a given peak is relative to the surrounding waveform. Specifically, prominence was calculated by integrating the difference between the height of the putative peak and the value of the local LFP waveform from *t*_*min*_ /2 before to *t*_*min*_ /2 after the time of the putative peak. To determine putative peaks, the function findpeaks was used as above, unless peak-like artifacts occurred close to other local maxima in the LFP. In these cases, we noticed that peaks exhibited a particular spectral signature (i.e., their occurrence was accompanied by increases in band power in a specific band, usually *δ* or *θ*). Thus, in these cases, we used band-limited power in conjunction with the LFP to identify putative peaks as follows. First, the function findpeaks was used to find maxima in the appropriate band power time series (with the relevant band determined by visual inspection) separated by a time *t*_*min*_. Second, putative peaks were identified as the local LFP maxima located no further in time than *t*_*jitter*_ from these maxima in band power. The time *t*_*jitter*_ was again determined by visual inspection, and was always smaller than the time *t*_*min*_ /2 (typically on the order of 10 ms).

### Eliminating Effects of Broadband LFP Deflections

After peaks were identified, the spectral parameters possibly affected by these peaks were excluded from both wavelet transforms and band power time series for all subsequent analyses, as follows. The number of data points removed from each time series was proportional to the inverse of the frequency represented by the time series, as seen in Figure 1c. Our intention, for each wavelet component and for each peak, was to identify data points at which the magnitude of the Morlet wavelet centered at the peak was greater than 10% of the maximum magnitude of the Morlet wavelet. For example, this would remove all timepoints within 1025 ms of a peak in the 1 Hz time series, all timepoints within 527 ms of a peak in the 2 Hz time series, and all timepoints within 35 ms of a peak in the 200 Hz time series. To implement this, a zero-one vector of peak timepoints (containing ones at each peak and zeros otherwise, so each peak is represented by a one) was convolved with the absolute values of the Morlet wavelets used for spectral analysis described above. The resulting time series were divided by their maxima, and time points at which these normalized time series were greater than 0.1 were set to NaN in all wavelet transforms. The results of these convolutions were also summed over the 6 frequency bands given above; these time series were divided by their maxima, and time points at which the normalized time series were greater than 0.1 were set to NaN in the band power time series. This method successfully removed most of the spectral contamination resulting from peak-like deflections in the LFP.

### Duration of Increases in Band-Limited Power

To determine the duration of increases in band power following carbachol infusion, we compared power within sliding 5 min windows (of 300 observations) to power during the pre-infusion baseline period, with one-sided unpaired t-tests at significance level p=0.05. These sliding windows were shifted with a step size of 30 s. Each timepoint was considered to be a timepoint at which *β* was increased, if it belonged to one sliding window in which power significantly increased from baseline.

### Identification of Snapshots of Oscillatory Dynamics Before and After Carbachol Infusion

The duration of recordings following infusion was longer than the duration of recordings before infusion, possibly biasing our calculation of increases in band-limited power (above). To account for the different duration of recordings pre- and post-infusion, we restricted our analysis of post-infusion recordings to the ten minutes surrounding the average of the midpoint of *β* increases in our previously recorded carbachol infusion data [McCarthy et al. (2011)]. This midpoint was 12 minutes, and so we analyzed only the recordings that took place 7 to 17 minutes post-infusion. Thus, the length of the analyzed pre- and post-infusion recordings was similar (7.9 ± 2.3 minutes pre-infusion and 10 minutes post-infusion).

Within this time period, the marked increases in *β* power visible in wavelet spectrograms (Fig. 2A) occurred at variable times, and could be preceded by decreases in *β* power [McCarthy et al. (2011)]. To account for this variability across animals, we compared “snapshots” of oscillatory dynamics – short periods characteristic of carbachol-induced increases in power at each frequency band – with periods of the same length, selected in the same way, from pre-infusion recordings. Since carbachol infusion induced power increases happened on a slow time scale of minutes, we used a 2.5 min long sliding window (with steps of 1 second) to identify candidate snapshots of increased oscillatory activity in the striatum. The criterion we used to select snapshots was a measure called “band power density” (BPD), designed to capture the burstiness of post-carbachol power increases, i.e. to indicate periods of especially dense but not continuous oscillatory activity.

**Figure 2:**
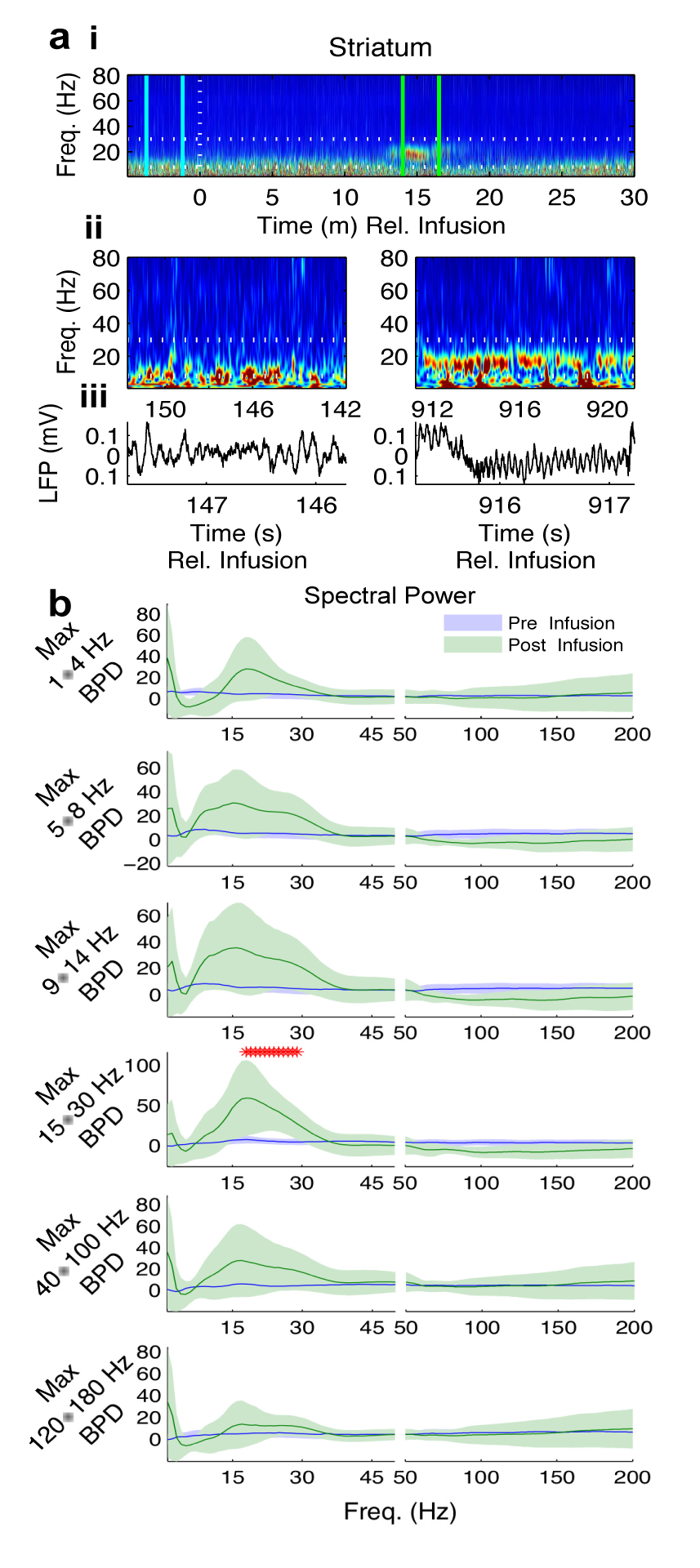
Carbachol selectively increases *β* oscillations in striatum. (A)Spectrograms of striatal LFP(representative animal).(i)The entire recording. White lines:horizontal,β band; vertical,time of infusion. Blue lines: 2.5 mins highest β BPD pre-infusion. Green lines: 2.5mins highest β BPD post-infusion. (ii)Tens in the middle of the period of highest β BPD pre-infusion(left)&post-infusion(right).(iii)Two s of LFP in the middle of the period of highest *β* BPD pre-infusion(left)&pots-infusion(right).(b)Average normalized striatal spectra for periods of highest BPD(mean±C.I.).Red stars: significant increases post-infusion.

**Figure 3:**
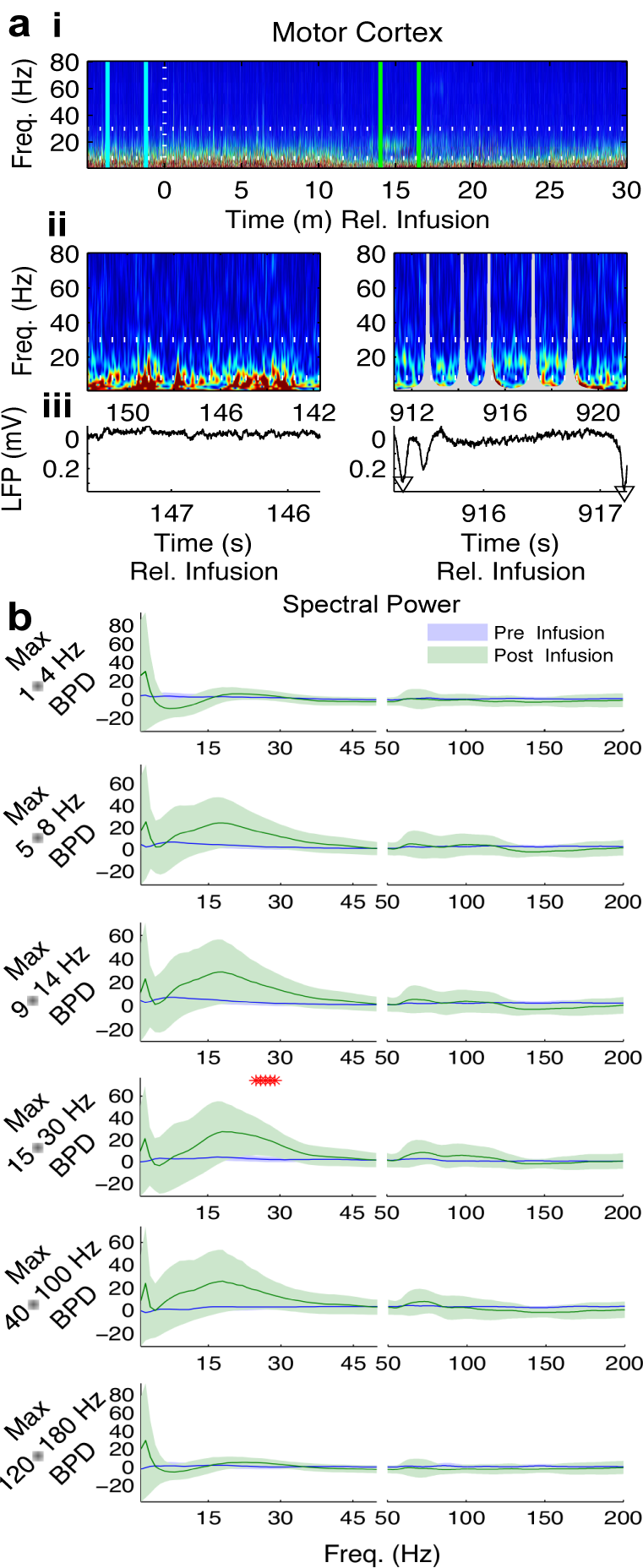
Carbachol selectively increases *β* oscillations in motor cortex. (A)Spectrograms of M1LFP(same representative animal as Fig.2a). (i)The entire recording. White lines: horizontal, β band; vertical, time of infusion. Blue lines: 2.5 mins highest β BPD pre-infusion. Green lines:2.5 mins highestβ BPD post-infusion. (ii)Tens in the middle of the period of highestβ BPD pre-infusion(left)& pos-infusion(right). (iii)Twos of LFP in the middle of the period of highest*β* BPD pre-infusion(left)&post-infusion(right). Overlays in Fig.3a ii and triangular symbols in Fig.3aiii(right panels) indicate identification and frequency-dependent removal of “peak”artifacts(see Methods, Fig.1).(b)Average normalized motor spectra for periods of highest BPD (mean± C.I.).Red stars:significant increases post-infusion.

To calculate BPD, we first identified putative timepoints of “high power” in each of the 6 frequency bands – *δ* (1–4 Hz), *θ* (5–8 Hz), *α* /low *β* (9–14 Hz), *β* (15–30 Hz), *γ* (40–100 Hz), and high-frequency oscillation (HFO, 120-180 Hz) –, as timepoints when the z-scored band power at that frequency was higher than 2 standard deviations from the mean over the pre-infusion period. To further eliminate contamination from peaks or smaller LFP deflections that were not removed by the peak detection algorithm described above, we eliminated putative high power timepoints occurring simultaneously with putative high power timepoints in any lower frequency band. The resulting high power timepoints were used to calculate BPD, as the number of high power timepoints per second.

Sliding windows were used to identify the 2.5 min period during which striatal BPD achieved the maximum over the time window between 7 and 17 minutes post-infusion. We used the same method to identify the 2.5 minutes of maximal striatal BPD over the entire pre-infusion period. Subsequent comparisons are made between the identified 2.5 minute snapshots before and after infusion.

### Statistical Comparisons of Pre- and Post-Infusion Power

To avoid fluctuations in lower frequency power from dominating our results (a possibility due to the power-law spectral signature of LFP data), and to allow comparisons between mice, we normalized each wavelet magnitude time series to the mean wavelet magnitude at that frequency during the pre-infusion period. Specifically, we normalized the wavelet magnitude time series as the percent change from mean pre-infusion wavelet magnitude.

Since wavelet transforms are computed using a convolution that essentially “slides” the wavelet time series along the LFP time series, the resulting estimates of power are not independent (i.e., adjacent time points influence each other, with adjacency determined by the length (in cycles) and the frequency of each wavelet – 1700 ms at 1 Hz, 850 ms at 2 Hz, and 60 ms at 200 Hz). To reduce the temporal correlation of the estimates of percent change in power, we divided the 2.5 minute periods analyzed into 150 non-overlapping 1 second long windows. One observation of normalized spectral power was calculated from each window by averaging the percent change in wavelet magnitude over that 1 second.

To determine statistical differences on individual animal level, we used the resulting 150 observations during the two periods, pre and post-infusion, by calculating paired t-tests between the two sets of 150 observations at each frequency. A t-test having a p-value below p = 0.01 for a frequency within the *β* band (15–30 Hz) indicated a significant change in *β* power pre- to post-infusion.

For population statistics across mice, average normalized spectral power over the pre- and post-infusion periods was calculated first for each mouse, and then the mean and standard error of normalized power were calculated across mice. We used a significance level of p = 0.05. Mean and 90% confidence intervals of normalized spectral power are plotted for the identified pre- and post-infusion periods, and paired one-sided t-tests across periods at each frequency were used to indicate a significant increase in power pre- to post-infusion, at level p = 0.05.

### Phase-Locking Between M1 and Striatum

**T**o examine phase-locking between **M**1 and striatum, we computed the magnitude of the mean resultant vector (MRV) of instantaneous phase differences (striatum – M1) for each frequency, for each second, within the identified 2.5 minute analysis window when striatal *β* BPD was highest, during pre- and post-infusion. First, the instantaneous phase of the wavelet transform in M1 was subtracted from the instantaneous phase of the wavelet transform in striatum, at each frequency and time. The circular mean phase difference at each frequency was then calculated for each second. The magnitude of the MRV provides a measure of the degree of phase consistency or phase-locking between the two recording sites at a particular frequency (phase-locking value, or PLV). As with spectral power, PLV was first normalized as a percentage of baseline (i.e. mean pre-infusion) PLV to allow comparisons across mice.

150 PLV values per period were used to compute differences on an individual level, by computing paired t-tests between periods for a set of 150 observations (pre- and post-infusion) per frequency. Group averaged PLV was then computed first for each mouse (across these 150 observations, and then averaged across mice (n = 12 mice). Paired one-sided t-tests (at significance level p = .05) across periods were used to detect increases in normalized PLV using these 12 observations at each frequency, and are displayed in Fig. S2 alongside the group average and 90% confidence intervals.

### Cross-Spectral Power and Partial Directed Coherence

**T**o fit the autoregressive model required for the calculation of cross-spectral power and partial directed coherence (PDC), it was necessary to remove peak artifacts in a way preserving the continuity of the signal in the time domain (Fig. 1d). **T**hus, instead of removing timepoints around peak artifacts from the spectral data in a frequency dependent way, we removed peak artifacts from the time series, by subtracting a suitably scaled average peak waveform from each peak, as follows. First, the set of peak artifacts for each animal was clustered into subsets. **T**hen, a scaled version of the mean peak waveform of each subset was removed from each peak within that subset. Scaling was accomplished by projected the time series around the peak onto the mean peak waveform. The resulting mean peak waveform was windowed and linearly offset, to ensure that the endpoints of the peak-subtracted data matched the endpoints of the raw data, and no discontinuities were introduced by the subtraction.

Following time-domain subtraction of peak artifacts, the pre-infusion baseline periods and the 2.5 min time series during the periods of highest BPD before and after carbachol infusion were divided into 10 s epochs. For each animal, each of these epochs was fit with an autoregressive model, using the MVGC Multivariate Granger Causality Toolbox [Barnett and Seth (2014)]. These autoregressive models were transformed into frequency-domain cross-spectral and PDC values, also using the MVGC Toolbox. The absolute values of cross-spectral and PDC measures during the 2.5 minutes of highest BPD were normalized by the mean values of these measures during the baseline period for each animal. Normalizations were calculated as the percent change from the baseline mean. At each frequency and for each period (pre- and post-infusion), fifteen normalized values of cross-spectral power and PDC resulted from the 10 s epochs obtained from the period of highest BPD. These 15 values were compared between pre- and post-infusion periods to obtain significant individual-level changes in cross-spectral power and PDC, at significance level p = 0.01. **T**he 12 sets of averaged normalized values (one from each animal) at each frequency pre- and post-infusion were compared with paired t-tests at significance level p = 0.05 to obtain group-level changes. The group-level difference between post- and pre-infusion cross-spectral power and PDC is plotted in Figure S2.

### Phase-Amplitude Coupling

For the calculation of phase-amplitude coupling, we removed peak artifacts in the same way as described above for the fitting of autoregressive models (Fig. 1d), and divided pre- and post-infusion periods of highest BPD into 10 s epochs. PAC comodulograms were computed for each epoch, 10 s as follows. For each pair of high-frequency amplitude and low-frequency phase time series (obtained using the wavelet transform as described above), an inverse entropy index [Pittman-Polletta et al. (2014)] was computed. First, a phase-amplitude “distribution” was constructed by: (i) dividing the phase domain (0, 2*p*] into 20 evenly-spaced bins; (ii) averaging the high-frequency amplitudes co-occuring with phases lying in each bin; and (iii) dividing by the sum across bins of these average amplitudes, to yield an empirical marginal distribution of high-frequency amplitude by low-frequency phase. Next, the entropy of this distribution of amplitude by phase – a measure of uniformity across phase – was computed, and normalized to yield a quantity between zero and one. **T**he resulting inverse entropy measure (IE) quantifies the degree of phase-amplitude dependence. An IE of zero indicates no phase-amplitude dependence (a uniform distribution of phase with respect to amplitude). An IE of one is obtained when all the amplitude occurs in a single phase bin. In general, higher IE is awarded to distributions exhibiting a stronger dependence of phase on amplitude, including distributions exhibiting multiple peaks.

For a finite data series, measurements of cross-frequency coupling will always take finite nonzero values, even when no coupling is present in the data. These nonzero values depend in part on the bandwidth, center frequency, and other properties of the observed signals. To remove these non-coupling-related influences, we used surrogate data to estimate “background” values of PAC. **T**hese background PAC values were estimated using epoch-shuffled surrogate data: for each channel and each 2.5 min snapshot, all non-simultaneous pairings of 10 s epochs were used to compute non-simultaneous sets of high- and low-frequency time series. **T**hese non-simultaneous time series were used to calculate a distribution of surrogate IE values. This distribution was used to z-score the observed IE for each epoch (at each pair of frequencies), yielding a “normalized” measure of significant coupling.

### Statistical Comparisons of Subgroups S+ and S-

When comparing within and across subgroups S+ (n=5) and S-(n=7), we performed t-tests of the average normalized values of each measure (per subgroup and per period) at each frequency. We used a significance level of p = 0.05. Mean and 90% confidence intervals of normalized spectral power are plotted for the identified pre- and post-infusion periods, and paired one-sided t-tests across periods at each frequency were used to indicate a significant increase in power pre- to post-infusion, at level p = 0.05. For comparisons of PAC between subgroups, the median PAC across epochs was computed for each animal, and the 12 resulting sets of averaged z-scored values (one from each animal) at each frequency pair were compared between subgroups S+ and S-, with ranksum tests at significance level p = 0.05 (since PAC measures are not approximately normal, even after z-scoring).

## RESULTS

**T**o determine how activation of striatal cholinergic receptors affects oscillations in both striatum and the larger CBT loop, we simultaneously recorded LFPs from striatum and M1 while infusing carbachol into the striatum of normal mice (Fig. 2a). In the striatum, carbachol infusion induced frequent bursts of strongly elevated *β* oscillations, with each burst lasting for hundreds of milliseconds to seconds (Fig. 2a). Consistent with our previous study [McCarthy et al. (2011)], such bursting was never observed before carbachol infusion.

### Carbachol Infusion Selectively Increases Striatal and M1 *β* Power

Carbachol infusion generated long-lasting increases in oscillatory power, with increases in *β* power lasting longer than increases in power at other frequencies. **T**o quantify periods of increased band power, we compared 5-minute sliding windows of power to all power pre-infusion (“baseline power”) within 7 different frequency bands, *δ* (1–4 Hz), *θ* (4–8 Hz), *α* /low *β* (8–15 Hz), *β* (15–30 Hz), *γ* (30–100 Hz), and high-frequency oscillation (HFO, 120–180 Hz) (Fig. S1). **T**he average duration of increased power was largest for the *β* band. *β* power increased over at least one 5 minute period in 10 out of 12 animals, with periods of elevated striatal *β* power lasting 17.06 ± 5.70 minutes (mean ± s.d.). **T**hese periods began 0.57 ± 1.01 minutes (mean ± s.d.) after infusion. Furthermore, in 4 of these animals, *β* remained increased at the end of the recording, 20 minutes post-infusion. In three long recordings (lasting ≥ 40 minutes post-infusion), the increase in striatal *β* power lasted 21.15 ± 7.29 minutes. While *β* increases were longer than increases in other frequencies, this difference was not statistically significant. Increases at multiple frequencies may in part be due to longer observation times post-infusion.

To eliminate the effect of longer observation times post-infusion, we examined oscillatory dynamics during the ten minute interval around the time of peak *β* increase in our previous results, namely from **7** to **17** minutes. **T**o account for the variable times of expression of carbachol-induced increases in power across animals, we examined 2.5-minute “snapshots” of the oscillatory dynamics during both pre- and post-infusion periods, in each band. More specifically, we compared periods of peak “band power density” (BPD) pre- and post-carbachol. BPD (the percentage of time when oscillatory power within a specific frequency band is significantly higher than baseline, see Methods and Fig. S2) is a measure that quantifies one of the most salient characteristics of observed increases in band power – namely, an increase in the percentage of time that the band power is higher than “normal” (Fig. 2 & 3 a).

During the period of highest striatal *β* BPD **7** to **17** minutes post-carbachol, spectral power in striatum and motor cortex increased selectively at high *β* frequencies (Figs. 2 & 3 b, row 4) at the group level. (At the individual level, striatal *β* power increased in 8 animals, and M1 *β* power increased in 6 animals.) Carbachol induced no other group-level changes in spectral power during this period (Figs. 2 & 3 b, row 4). Carbachol also failed to induce increases in spectral power at any frequency during periods of highest BPD for non-*β* bands (Fig. 2 & 3 b). Interestingly, all periods of highest non-*β* BPD had distinct, though statistically non-significant, power peaks in the *β* frequency range. **T**hese results extend previous findings [McCarthy et al. (2011)] by showing that carbachol selectively increases *β* oscillations in the striatum and in M1.

### Increased Striatum → M1 PDC Accompanies Simultaneous Carbachol-Induced *β* Increases

In PD, elevation of *β* power is accompanied by changes in *β* phase coordination within the CBT loop [Stein and Bar-Gad (2013); Cagnan et al. (2015)]. To further explore whether carbachol-induced *β* increases were accompanied by changes in *β* coordination, we honed in on the subset of animals (n=5) showing simultaneous increases in *β* power in both M1 and striatum (subgroup S+). In this subgroup, both striatal and M1 *β* power were elevated in the frequency ranges 23–30 Hz and 23–31 Hz, respectively, at the subgroup level (Fig. 4).

**Figure 4:**
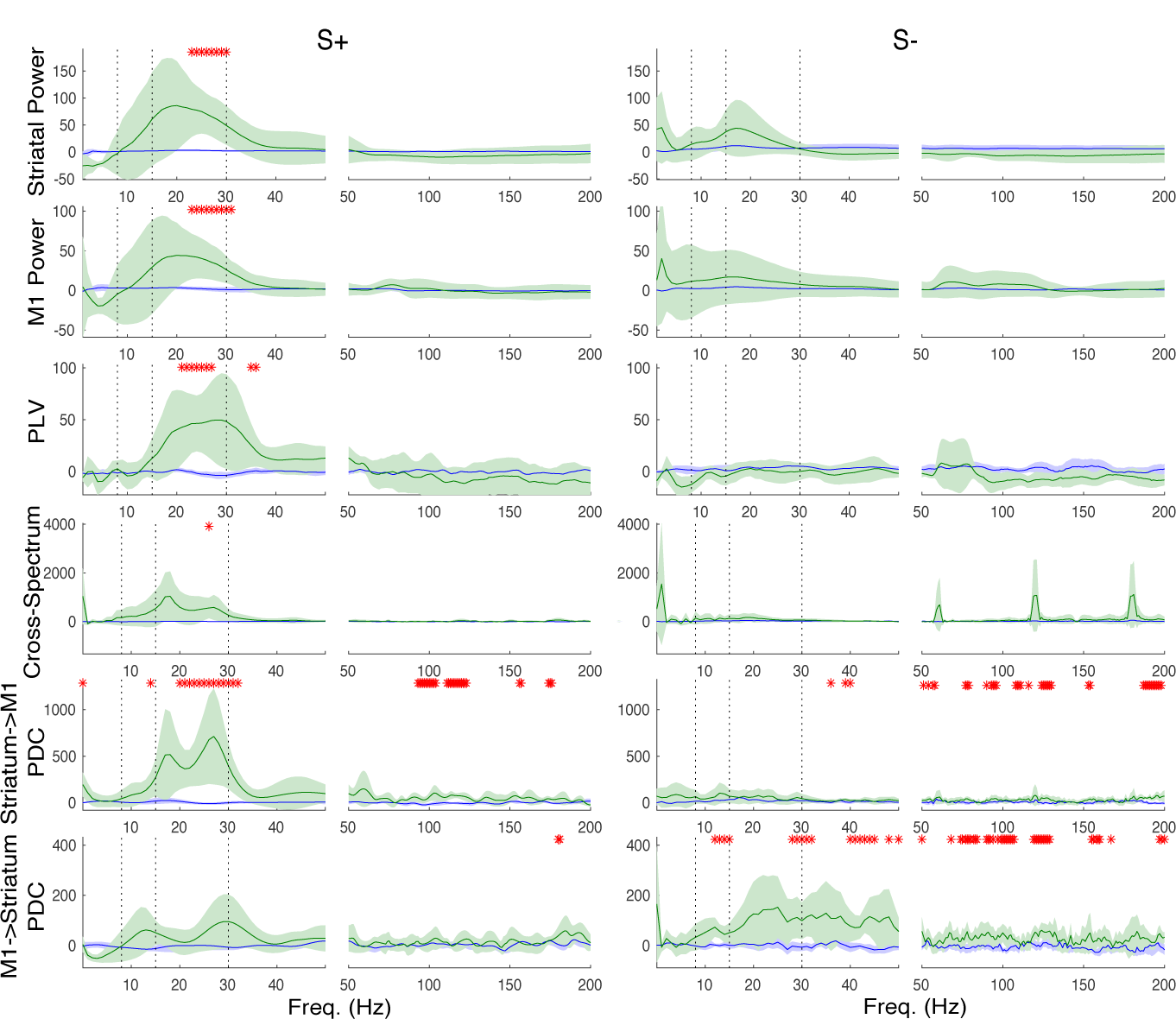
Simultaneous increases in striatal and motor *β* power and coordination are dichotomously expressed after striatal carbachol infusion. Average spectral power, PLV, cross spectrum, & partial directed coherence (mean ± C.I.) for subgroups S+ and S-. Red stars: significant increases post-infusion.

The data from subgroup S+ show that when activation of striatal cholinergic receptors elevates *β* oscillations in both striatum and M1, it also increases *β* coordination in corticostriatal circuits. **T**he phase locking value (PLV) between simultaneously recorded LFPs in striatum and M1 was robustly and selectively increased (during periods of highest striatal *β* BPD) in the higher *β* frequency range (21–27 Hz) in S+ (Fig. 4). We also examined the cross spectrum between M1 and striatum (another measure of rhythmic coordination) and found an increase at 26 Hz in S+ (Fig. 4).

**T**o characterize the directionality of elevations in *β* coordination in S+, we measured partial directed coherence (PDC) between striatal and M1 LFPs during the period of highest striatal BPD. PDC is a measure of how (frequency-specific) fluctuations in one time series predict (frequency-specific) fluctuations in another time series, indicating the flow of information within a given frequency band between structures [Barnett and Seth (2014)]. In subgroup S+, PDC analyses revealed that carbachol infusion increased the influence of the striatum on M1 in the *β* frequency range from approximately 20–32 Hz, and also the influence of striatal *γ* (93–104 & 111–123 Hz) band activity on M1 (Fig. 4). In contrast, PDC from M1 to striatum did not increase almost at all in S+, except an increase at 180–181 Hz (Fig. 4).

### S-was Distinguished by Increased M1→Striatum PDC and *β*-HFO Phase-Amplitude Coupling

We next sought to determine the possible causes of the lack of simultaneous increases in *β* power in the remaining 7 individuals (subgroup S-). Within this subgroup, no increases in *β* power or *β* coordination were observed at the subgroup level. However, increases in M1→striatum *β* PDC were observed (from 12–15 and 28–32 Hz), while *β* PDC from striatum to M1 did not increase. Additional increases in PDC were observed in both directions in the *γ* band in S-(Fig. 4).

**T**o test whether these animals might be asleep, we compared the *δ*/*θ* ratio (a measure increased during deep sleep) during the baseline period between subgroups, finding no difference (Fig. S2). In a further search for markers of differences in dynamic states between the two subgroups, we examined cross-frequency coupling. Cross-frequency coupling between brain rhythms within the CB**T** loop is implicated in the function (and dysfunction) of the motor system, and changes in cross-frequency coupling may characterize behaviorally relevant changes in CB**T** dynamics [Lopez-Azcarate et al. (2010); López-Azcárate et al. (2013); von Nicolai et al. (2014)]. We examined phase-amplitude coupling (PAC), in which the amplitude of a high frequency oscillation exhibits a statistical dependence on the phase of a low frequency oscillation [Canolty and Knight (2010); Pittman-Polletta et al. (2014)], in both striatal and M1 LFPs, pre- and post-infusion, in subgroups S+ and S-. Subgroup S-showed *β*-HFO PAC that was clearly absent in subgroup S+ (Fig. 5). This *β* phase modulation of HFO power was present during both pre- and post-infusion periods of highest striatal *β* BPD. Furthermore, subgroup S+ showed increased *α*-*γ* PAC relative to subgroup S-post-carbachol (Fig. 5). This result gives more evidence that subgroups S+ and S-are in distinct dynamical states, even before carbachol infusion occurs. (No clear changes emerged when comparing pre- and post-infusion PAC in either subgroup.)

**Figure 5:**
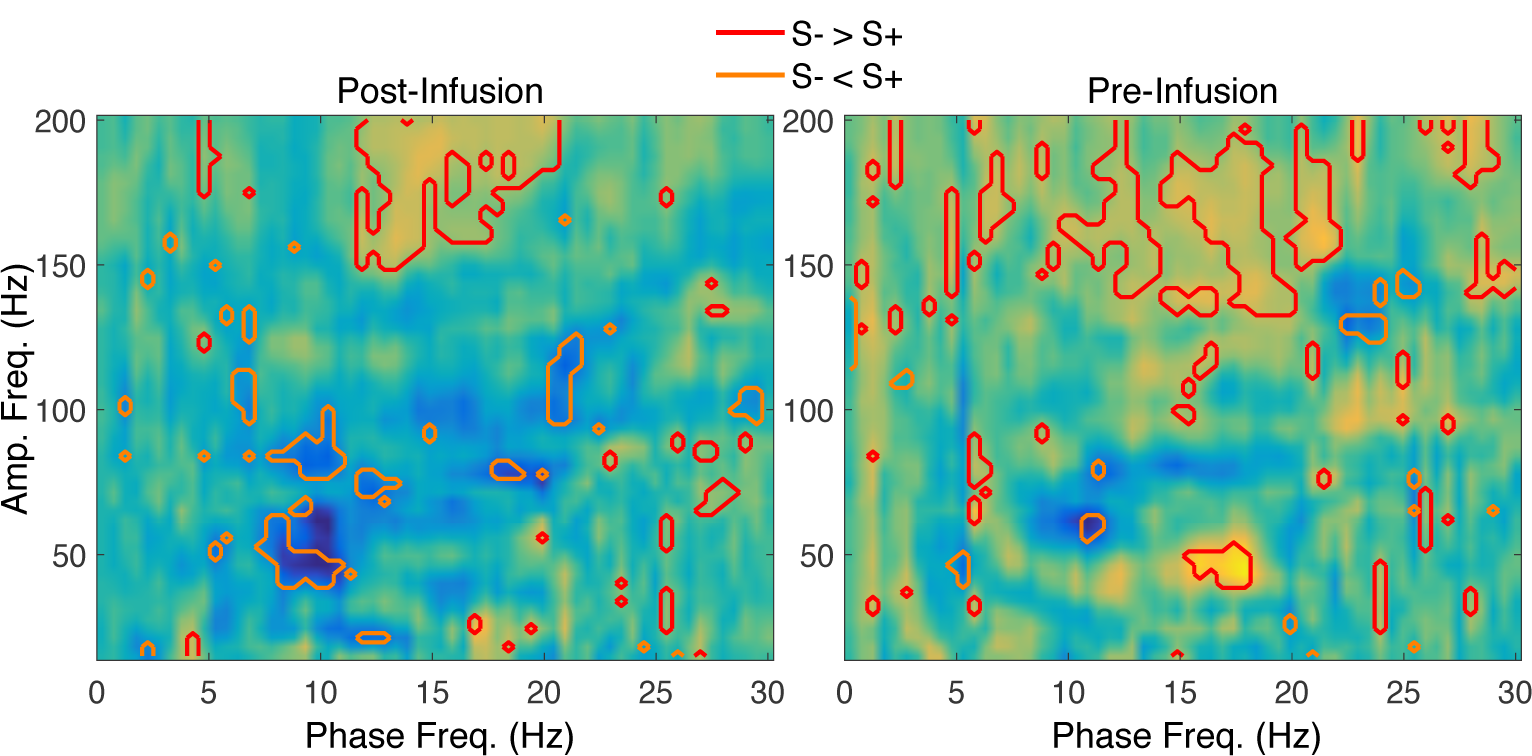
M1 phase-amplitude coupling distinguishes subgroups S+ and S-. The difference in median PAC between subgroups S+ and S-(S+ subtracted from S-). Red contours: significantly increased PAC in subgroup S-as compared to subgroup S+.

### The Directionality of PDC Increases was Frequency-Specific

Examining the frequencies at which individual animals exhibited changes in PDC also suggested an association between information flow and *β* subbands (Fig. 6). Most animals exhibited increases in striatal to M1 PDC at frequencies between 11–16 and 20–26 Hz. In contrast, most animals exhibited increases in M1 to striatal PDC at frequencies above 29 Hz. **T**his suggests that distinct *β* subbands are differentially modulated following striatal carbachol infusion.

**Figure 6:**
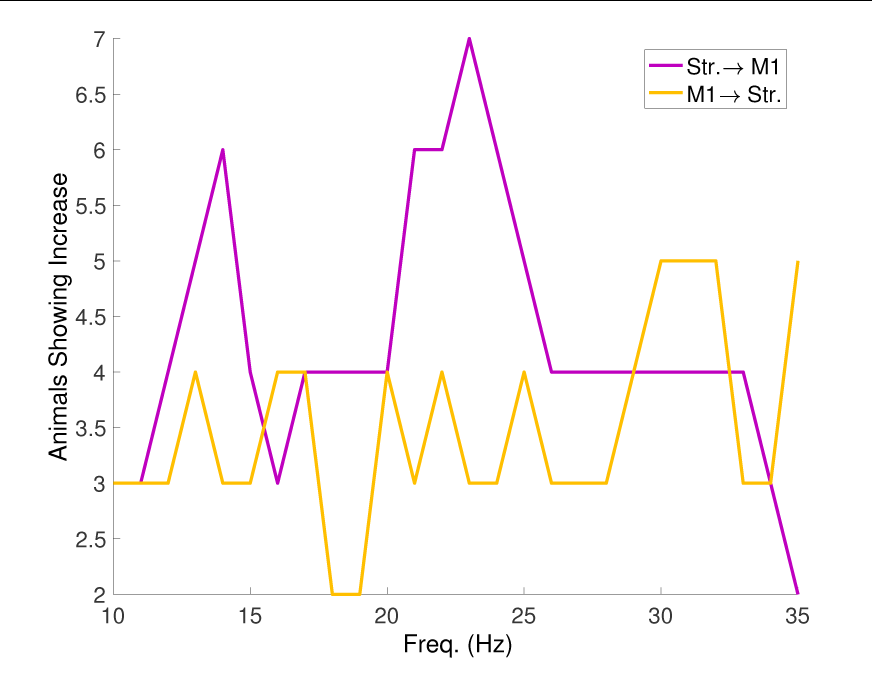
The direction of *β* PDC increases is frequency-dependent. The number of animals showing carbachol-induced increases in PDC at each frequency between 10 and 35 Hz.

## DISCUSSION

Infusion of carbachol into the striatum of normal mice selectively elevated *β* oscillations in both striatum and M1 and brought to light two independent dynamical states in CBT circuits. The first state (S+) was defined by elevated *β* frequency oscillations in both striatum and M1, and characterized by: (1) increased coordination between *β* oscillations in these two structures as measured by PLV and cross-spectral power; and (2) increased information flow from striatum to M1 in the *β* frequency band. In the second state (S-), power in all frequency bands, including *β*, remained at pre-infusion levels in both striatum and M1. Of note, *β*-HFO PAC was robustly present in S-(and increased relative to S+) both pre- and post-infusion. This finding strongly suggests S-is a manifestation of a different CBT dynamical state than S+ and that the difference existed prior to carbachol infusion. No changes from baseline in PLV or cross-spectral power were noted in state S-, however, this state also showed increased information flow from M1 to striatum in the *β* frequency range, further distinguishing S- and S+ as distinct dynamical states. Indeed, such state dependence may explain inconsistent findings on acute *β* increases in frontal cortex: experiments on systemic dopamine depletion in anesthetized rats, which failed to increase *β* power in ECoG recorded from somatomotor cortex [Mallet et al. (2008b)], may be observations of our state S-; while experiments in which acute striatal dopamine depletion increased *β* power in M1 in awake DAT knockout (DAT-KO) mice [Costa et al. (2006)] may be observations of our state S+.

In contrast to recent experimental work showing that optogenetic stimulation of striatal cholinergic interneurons (sChIs) induces broadband increases in oscillatory power in striatum and M1 [Kondabolu et al. (2016)], the current study shows a selective elevation of power in the *β* frequency range within corticostriatal circuits. These increases can be attributed to the stimulation of striatal cholinergic receptors: In our previous study [McCarthy et al. (2011)], control experiments showed that the changes in beta **LFP** in striatum after carbachol infusion were not due to the pressure induced by drug infusion, which can activate mechano-sensitive mechanisms in striatum, since infusion of a lower concentration of carbachol (0.1 - 0.2 mM infused with up to 5 *μ*L) did not change striatal beta power throughout the entire infusion period. The carbachol-induced rise in beta is likely mediated through muscarinic receptors, since our previous study using optogenetic stimlation of striatal cholinergic interneurons showed beta oscillations remained elevated in the presence of the nicotinic antagonist mecamylamine, but were diminished when muscarinic receptors were blocked by scopolamine [Kondabolu et al. (2016)]. In subgroup S+, all significant increases in power and cross-spectral power, both in striatum and M1, occurred within the **21 - 31 Hz** frequency range. **PLV** was also largely constrained to the *β* frequency range in S+. In contrast, optogenetic stimulation of sChIs increases *α, β*, and low and high *γ* power in striatum, and *β* and low and high *γ* power in M1 [Kondabolu et al. (2016)]. What could be the source of the sChI-induced elevations in *α*, low *γ*, and high *γ* power, absent in our subgroup S+? The previous study suggests that sChI-induced high *γ* is not attributable to activation of striatal cholinergic receptors, since sChI-induced high *γ* was not reduced by either muscarinic or nicotinic blockade [Kondabolu et al. (2016)]. Since sChIs co-release glutamate with **AC**h [Higley et al. (2011)], perhaps glutamatergic mechanisms are involved in producing striatal high *γ* oscillations. Alterations in *α* oscillations within cortico-striatal circuits, however, may be due to cholinergic mechanisms. Indeed, although *α* power was not elevated in striatum in either S+ or S-, alpha PDC was significantly increased in both subgroups, although in opposite directions, indicating that activation of striatal cholinergic receptors can modulate *α*-band information transfer in a state-dependent manner. **T**he source of elevated sChI-induced low *γ* is unclear, since sChI-induced low *γ* was diminished following muscarinic blockade [Kondabolu et al. (2016)]. Perhaps the discrepancy between the current pharmacologic and the previous optogenetic study stems from the lower frequency resolution resulting from shorter trial lengths in the previous study.

The PDC during periods of elevated *β* in S+ suggests that the flow of information in the elevated *β* band is directed from striatum to M1. However, interestingly, in S-directed information flow increases from M1 to striatum even though *β* power remains at baseline levels in both structures. These dichotomous results for PDC may help explain results on rats rendered parkinsonian by 6-OHDA lesion, which show bidirectional PDC between cortex and striatum in the *β* frequency range with the flow of information predominantly from striatum to cortex [Belic et al. (2016)]. Our results indicate that in states with high striatal cholinergic tone such as occurs in the 6-OHDA rat [Ikarashi et al. (1997)], PDC is increased between striatum and cortex, with the direction of increased information flow depending on the dynamical state expressed in CBT circuits. Of note, 3 of the 7 animals in subgroup S-show an increase in striatal *β* power in the absence of a corresponding increase in M1 *β* power, while only 1 animal shows an increase in M1 *β* power in the absence of striatal *β* increases. Like the subgroup specific increases in *β* PDC that we observed, this result provides evidence for the independence of striatal and M1 *β* oscillations, and shows that it is possible to increase striatal *β* power independently from M1 *β* power. The independence of M1 and striatal *β* generators is also supported by our results showing that more mice exhibited increases in M1→striatum PDC at frequencies above 29 Hz, whereas more mice showed striatum→M1 PDC at lower *β* frequencies (11 - 16 Hz and 20 - 26 Hz). These frequency- and state-dependent increases in *β* PDC are consistent with results that suggest M1 as a generator of high *β* frequencies [Yamawaki et al. (2008); Lacey et al. (2014)], and striatum as a generator of a range of *β* frequencies [McCarthy et al. (2011)]. Notably, in S-, M1 PAC between *β* and HFOs only occurs for low *β* frequencies (less than approximately 20 Hz), further evidence for the existence of at least two frequency ranges within the *β* band having different properties and perhaps different mechanistic sources. This idea is consistent with evidence that low and high *β* are distinct entities in the parkinsonian basal ganglia, responding differently to dopaminergic therapy and DBS, and potentially correlating with different motor related behavior and motor symptoms in PD [Stein and Bar-Gad (2013); Lopez-Azcarate et al. (2010); Toledo et al. (2014); Priori et al. (2004); Foffani et al. (2005); Blumenfeld et al. (2017); Oswal et al. (2016); Marceglia et al. (2009)]. Our work suggests that which of these potential *β*-generating mechanisms predominates at any given time may depend on the intrinsic state of the CBT loop.

Importantly, carbachol infusion did not increase *β* power in either region in 3 of 12 animals, or at the subgroup level in S-. This suggests that there are dynamic or constitutive factors that may prevent the appearance of exaggerated *β* oscillations in the CBT network. These factors may play a role in conferring resistance to the negative symptoms of movement disorders. Our results provide evidence that S-is a naturally-occuring network state preventing the elevation and coordination of *β* in CBT circuits (and perhaps its associated motor pathology). Further characterizing the cellular and network mechanisms underlying this state may have therapeutic value in treating the motor symptoms associated with elevated *β* oscillations in Parkinson’s disease patients. One of the most striking differences between states S+ and S-is the elevation of M1 low-beta/HFO PAC in S-. Although both the mechanisms of generation and function of HFOs are unknown, it is interesting to note that the HFO frequency that couples to low *β* (130–200 Hz) is similar to the range of effective frequencies used for deep brain stimulation (DBS) (around 130–185 Hz) for the alleviation of motor symptoms in PD [Kuncel and Grill (2004)]. Perhaps DBS amplifies intrinsic M1 mechanisms that prevent the propagation of elevated *β* oscillations around the CBT loop. Indeed, HFOs also appear in M1 after severe dopamine depletion in rats, with the amount of *β* correlating inversely with the amount of HFOs [Brys et al. (2017)], consistent with the idea that the HFOs may offer some protective effect against exaggerated *β*. Further research into the mechanisms and functions of these HFOs is needed to shed light on these possibilities. Of additional note, the low *β*-HFO PAC is high both before and after carbachol infusion in state S-, which not only lends additional evidence to the idea that S- and S+ represent two naturally-occuring and functionally distinct states of CBT dynamics, but also suggests that the dynamical state prior to perturbation by carbachol may be a critical regulator of *β* propagation in CBT circuits. Further characterizing the dynamics present during state S-may provide additional insight into a normal CBT network state that is capable of preventing the propagation of *β* oscillations throughout the CBT loop. Finally, although we did not monitor behavior in these experiments, our findings suggest that the difference in states cannot be accounted for by deep sleep versus wakefulness as there was no difference in the *δ*/*θ* ratio between these subgroups (Fig. S1).

The present study shows that *β* can be acutely elevated in striatum and M1 after stimulation of striatal cholinergic receptors, showing that plastic changes due to chronic loss of dopamine are not required to produce robust *β* in CBT circuits. Our previous computational modeling work suggests that networks of striatal medium spiny neurons (MSNs) can robustly and reversibly produce *β* frequency oscillations in response to perturbations that increases the spiking rate of the MSNs, including heightened striatal cholinergic tone and lowered dopaminergic tone [McCarthy et al. (2011)]. The experimental results reported here support this model of *β* generation in basal ganglia. Additional support for this model comes from studies showing elevated *β* oscillations in striatum and M1 after acute loss of dopamine in DAT-KO mice concurrently with parkinsonian motor symptoms [Fuentes et al. (2009); Costa et al. (2006)], suggesting PD *β* pathology and its associated symptoms may not require long-term plastic changes to basal ganglia structures.

The striatal *β* model [McCarthy et al. (2011)] introduces a mechanism capable of producing exaggerated *β* oscillations acutely. These acute changes in *β* may coexist with *β*-producing mechanisms dependent on slow, plastic changes consequent to dopamine depletion, as posited by proposed models of *β* generation in Parkinson’s disease that are dependent on oscillatory dynamics in STN/GPe and GPe/striatal circuits [Holgado et al. (2010); Corbit et al. (2016); Shouno et al. (2017); Terman et al. (2002); Rubin and Terman (2004)]. Similar STN/GPe models may be capable of producing acute increases in *β* [Kumar et al. (2011)]. However, to our knowledge, there is no evidence for acute *β* increases in STN or GPe following dopamine depletion (or any other experimental manipulation). Indeed there is some evidence that acute *β* increases are missing in STN following DA depletion: in the 6-OHDA rat model of Parkinson’s disease, *β* oscillations are not elevated in the STN until several days after dopaminergic lesion, and the STN does not show increases in *β* oscillations after acute disruption of D1 or D2 receptor signaling [Mallet et al. (2008b)], consistent with the necessity of plastic changes for the production of STN *β* oscillations. We note that it remains unclear how cholinergic tone affects dopamine in striatum during carbachol infusion [Rice et al. (2011); Threlfell et al. (2012)], and the carbachol-induced *β* may be working by acutely decreasing dopamine in striatum. However, although recordings from STN were not obtained, it is unlikely that the plastic changes to the STN/GPe circuits due to chronic loss of dopamine which appear necessary for elevated STN *β* were achieved during the relatively short duration of post-infusion recordings in this study.

A result commonly claimed to refute the possibility of striatal *β* generation in PD comes from experiments done on MPTP monkeys [Tachibana et al. (2011)], in which local blockade of projections from striatum to a microregion of GPe failed to abolish *β* rhythms in that microregion. However, the small proportion of striato-pallidal synapses blocked in these experiments leaves open the possibility that *β* oscillations propagated from striatum to the blocked portion of GPe through the intact (majority of) projections and pathways connecting these two regions. In particular, while the projections from striatum to GPe are convergent, the loop from STN to GPe and back is highly divergent [Haber (2010)], and may have relayed *β* from the intact surrounding GPe to the blocked microregion.

Antimuscarinics were a mainstay of therapy for Parkinson’s disease until the advent of L-dopa in the 1960s [Pisani et al. (2007)] and the current study suggests that the main site of their efficacy may reside in their ability to lower striatal cholinergic tone. However, although striatal neurons may experience hypercholinergia, other cholinergic centers in the brain degenerate in PD, including the basal forebrain cholinergic system, which is the primary source of cholinergic input to the cortex[Bohnen and Albin (2011)]. Thus, although PD patients can experience some relief of symptoms from anticholinergics, the additional hypocholinergia produced in non-striatal sites by systemic anticholinergics in PD patients likely contribute to the limiting side effects of anticholinergic drugs, including cognitive impairment, confusion, memory loss, and nausea [Fox (2013); Langmead et al. (2008); Lees (2005)]. In contrast, targeted striatal anticholinergic interventions alleviate parkinsonian motor symptoms in 6-OHDA mice [Maurice et al. (2015); Ztaou et al. (2016)], suggesting that the therapeutic effect of systemic anticholinergics may be due to the lowering of cholinergic tone in striatum. The present study suggests that anticholinergics may exert their beneficial effects in PD by reversing the network dynamics producing exaggerated *β* oscillations within CBT circuits.

In the present study, we find that activation of striatal cholinergic receptors can selectively elevate the power and coordination of *β* frequency oscillations in striatum and M1. Elevation of striatum and M1 *β* oscillations by increased striatal cholinergic tone depends on the dynamic state of the CBT loop prior to and during carbachol infusion. Our results demonstrate a prominent role for striatal cholinergic receptors as state-dependent modulators of *β* frequency oscillations throughout the CBT loop, and have widespread implications for the importance of these receptors in the control of voluntary movement and its loss in PD.

## ACKNOWLEDGMENTS

X.H. acknowledges funding from NIH Director’s New Innovator Award 1DP2NS082126, NINDS Grant 1R21NS078660, Pew Foundation, Alfred P. Sloan Foundation, Boston University Biomedical Engineering Department, and Boston University Photonic Center. X.H. and M.M.M acknowledge CRCNS NIH Grant 1R01NS081716. N.K. acknowledges NSF DMS-1042134.

## AUTHOR CONTRIBUTIONS

A.Q, N.K., X.H., and M.M.M. designed the study. N.K., M.M.M., and X.H. supervised the study. B.R.P.P. designed and performed the data analysis. A.Q. and A.I.M. performed the experiments and collected data.

M.R. assisted with data analysis. K.K. assisted with data collection. B.R.P.P., N.K., X.H., and M.M.M. wrote the manuscript.

## DATA ACCESSIBILITY

All code used to perform the data analysis can be accessed online at https://github.com/benpolletta/PD_Data.

---

**Figure S1:**
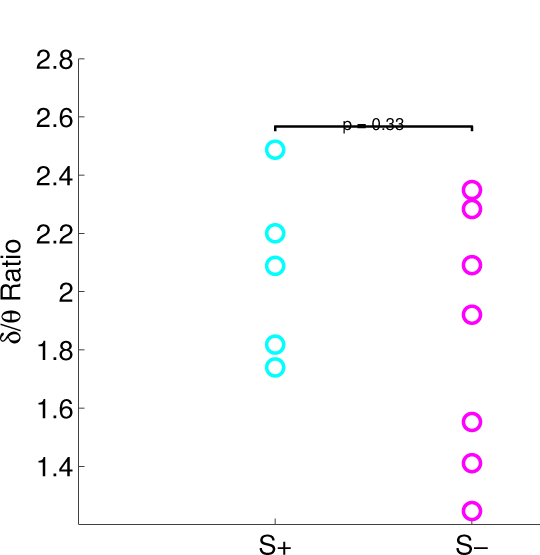
*δ*/*θ* ratio does not distinguish subgroups S+ and S-. The mean *δ*/*θ* ratio during baseline for animals in subgroups S+ and S-.

**Figure S2:**
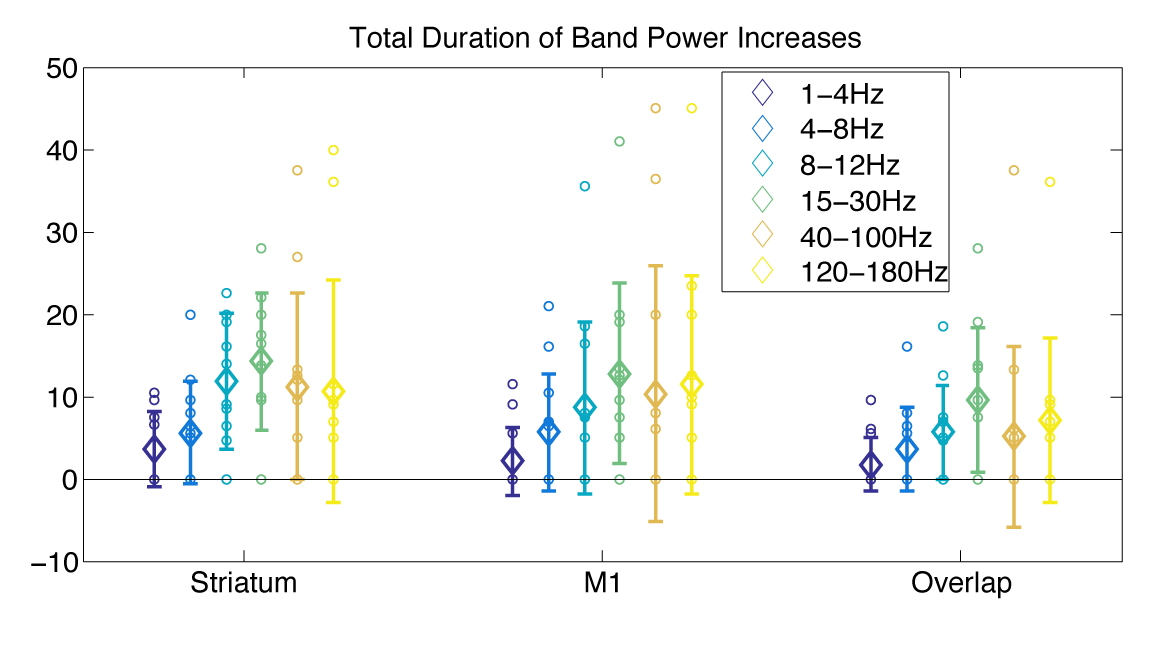
Band power increases relative to baseline in all bands, with *β* power increases having the longest duration. Duration of band power increases (mean ± S.E.) is shown for all frequency bands, for the striatal LFP (left), the M1 LFP (middle), and for both LFPs (right).

**Figure S3:**
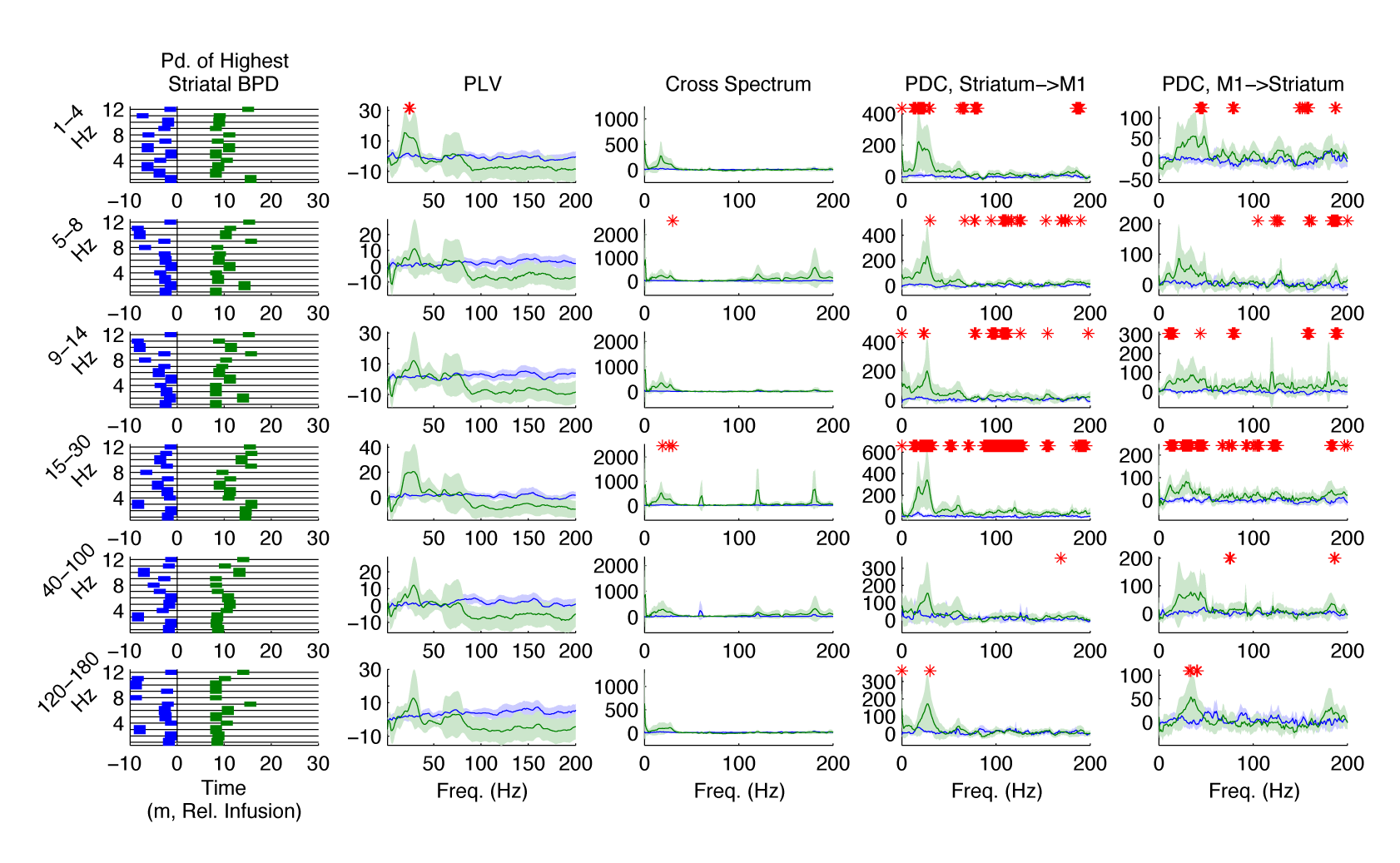
Periods of highest BPD and *β* coordination. Column 1: The 2.5 min time periods of highest band power density (BPD) for all experiments. Blue bars: period of highest BPD pre-infusion. Green bars: period of highest BPD post-infusion. Thick bars: subgroup S+. Thin bars: subgroup S-. Black vertical line: infusion begins. Column 2: Average normalized PLV for periods of highest BPD (mean ± C.I.). Red stars: significant increases post-infusion.Column 3: Average normalized cross spectrum for periods of highest BPD (mean ± C.I.). Red stars: significant increases post-infusion. Column 4: Average normalized PDC (striatum→M1) for periods of highest BPD (mean ± C.I.). Red stars: significant increases post-infusion. Column 5: Average normalized PDC (M1→striatum) for periods of highest BPD (mean ± C.I.). Red stars: significant increases post-infusion.

## Abbreviations

ACh: acetylcholine
BPD: band power density
CBT: cortex-basal ganglia
DAT: dopamine transporter
GPe: globus pallidus externa
HFOs: high frequency oscillations
KO: knockout
LFP: local field potential
M1: primary motor cortex
PAC: phaseamplitude coupling
PD: Parkinsons’
disease: 
PDC: partial directed coherence
PLV: phase-locking value
sChI: striatal cholinergic interneuron
STN: subthalamic nucleus

## REFERENCES

Barnett, L. and Seth, A. K. (2014) The mvgc multivariate granger causality toolbox: a new approach to granger-causal inference. Journal of neuroscience methods, 223, 50–68.

Belic, J. J., Halje, P., Richter, U., Petersson, P. and Hellgren Kotaleski, J. (2016) Untangling cortico-striatal connectivity and cross-frequency coupling in l-dopa-induced dyskinesia. Front Syst Neurosci, 10, 26.

Blumenfeld, Z., Koop, M. M., Prieto, T. E., Shreve, L. A., Velisar, A., Quinn, E. J., Trager, M. H. and Bronte-Stewart, H. (2017) Sixty-hertz stimulation improves bradykinesia and amplifies subthalamic low-frequency oscillations. Mov Disord, 32, 80–88.

Bohnen, N. I. and Albin, R. L. (2011) The cholinergic system and parkinson disease. Behav Brain Res, 221, 564–73.

Brys, I., Nunes, J. and Fuentes, R. (2017) Motor deficits and beta oscillations are dissociable in an alpha-synuclein model of parkinson’s disease. Eur J Neurosci.

Cagnan, H., Duff, E. P. and Brown, P. (2015) The relative phases of basal ganglia activities dynamically shape effective connectivity in parkinson’s disease. Brain, 138, 1667–1678.

Canolty, R. T. and Knight, R. T. (2010) The functional role of cross-frequency coupling. Trends in Cognitive Sciences, 14, 506–515.

Corbit, V. L., Whalen, T. C., Zitelli, K. T., Crilly, S. Y., Rubin, J. E. and Gittis, A. H. (2016) Pallidostriatal projections promote beta oscillations in a dopamine-depleted biophysical network model. J Neurosci, 36, 5556–71.

Costa, R. M., Lin, S. C., Sotnikova, T. D., Cyr, M., Gainetdinov, R. R., Caron, M. G. and Nicolelis, M. A. (2006) Rapid alterations in corticostriatal ensemble coordination during acute dopamine-dependent motor dysfunction. Neuron, 52, 359–69.

Ding, J., Guzman, J. N., Tkatch, T., Chen, S., Goldberg, J. A., Ebert, P. J., Levitt, P., Wilson, C. J., Hamm, H. E. and Surmeier, D. J. (2006) Rgs4-dependent attenuation of m4 autoreceptor function in striatal cholinergic interneurons following dopamine depletion. Nat Neurosci, 9, 832–42.

Foffani, G., Bianchi, A. M., Baselli, G. and Priori, A. (2005) Movement-related frequency modulation of beta oscillatory activity in the human subthalamic nucleus. J Physiol, 568, 699–711.

Fox, S. H. (2013) Non-dopaminergic treatments for motor control in parkinson’s disease. Drugs, 73, 1405–15.

Fuentes, R., Petersson, P., Siesser, W. B., Caron, M. G. and Nicolelis, M. A. (2009) Spinal cord stimulation restores locomotion in animal models of parkinson’s disease. Science, 323, 1578–82.

Haber, S. N. (2010) Integrative Networks Across Basal Ganglia Circuits-Chapter 24. Elsevier Science & Technology.

Higley, M. J., Gittis, A. H., Oldenburg, I. A., Balthasar, N., Seal, R. P., Edwards, R. H., Lowell, B. B., Kreitzer, A. C. and Sabatini, B. L. (2011) Cholinergic interneurons mediate fast vglut3-dependent glutamatergic transmission in the striatum. PLoS One, 6, e19155.

Holgado, A. J., Terry, J. R. and Bogacz, R. (2010) Conditions for the generation of beta oscillations in the subthalamic nucleus-globus pallidus network. J Neurosci, 30, 12340–52.

Ikarashi, Y., Takahashi, A., Ishimaru, H., Arai, T. and Maruyama, Y. (1997) Regulation of dopamine d1 and d2 receptors on striatal acetylcholine release in rats. Brain Res Bull, 43, 107–15.

Kharkwal, G., Brami-Cherrier, K., Lizardi-Ortiz, J. E., Nelson, A. B., Ramos, M., Del Barrio, D., Sulzer, D., Kreitzer, A. C. and Borrelli, E. (2016) Parkinsonism driven by antipsychotics originates from dopaminergic control of striatal cholinergic interneurons. Neuron, 91, 67–78.

Kondabolu, K., Roberts, E. A., Bucklin, M., McCarthy, M. M., Kopell, N. and Han, X. (2016) Striatal cholinergic interneurons generate beta and gamma oscillations in the corticostriatal circuit and produce motor deficits. Proc Natl Acad Sci U S A.

Kuhn, A. A., Kempf, F., Brucke, C., Gaynor Doyle, L., Martinez-Torres, I., Pogosyan, A., Trottenberg, T., Kupsch, A., Schneider, G. H., Hariz, M. I., Vandenberghe, W., Nuttin, B. and Brown, P. (2008) High-frequency stimulation of the subthalamic nucleus suppresses oscillatory beta activity in patients with parkinson’s disease in parallel with improvement in motor performance. J Neurosci, 28, 6165–73.

Kuhn, A. A., Kupsch, A., Schneider, G. H. and Brown, P. (2006) Reduction in subthalamic 8-35 hz oscillatory activity correlates with clinical improvement in parkinson’s disease. Eur J Neurosci, 23, 1956–60.

Kuhn, A. A., Tsui, A., Aziz, T., Ray, N., Brucke, C., Kupsch, A., Schneider, G. H. and Brown, P. (2009) Pathological synchronisation in the subthalamic nucleus of patients with parkinson’s disease relates to both bradykinesia and rigidity. Exp Neurol, 215, 380–7.

Kumar, A., Cardanobile, S., Rotter, S. and Aertsen, A. (2011) The role of inhibition in generating and controlling parkinson’s disease oscillations in the basal ganglia. Frontiers in Systems Neuroscience, 5, 86.

Kuncel, A. M. and Grill, W. M. (2004) Selection of stimulus parameters for deep brain stimulation. Clin Neurophysiol, 115, 2431–41.

Lacey, M. G., Gooding-Williams, G., Prokic, E. J., Yamawaki, N., Hall, S. D., Stanford, I. M. and Woodhall, G. L. (2014) Spike firing and ipsps in layer v pyramidal neurons during beta oscillations in rat primary motor cortex (m1) in vitro. PLoS One, 9, e85109.

Langmead, C. J., Watson, J. and Reavill, C. (2008) Muscarinic acetylcholine receptors as cns drug targets. Pharmacol Ther, 117, 232–43.

Lees, A. (2005) Alternatives to levodopa in the initial treatment of early parkinson’s disease. Drugs Aging, 22, 731–40.

López-Azcárate, J., Nicolás, M. J., Cordon, I., Alegre, M., Valencia, M. and Artieda, J. (2013) Delta-mediated cross-frequency coupling organizes oscillatory activity across the rat cortico-basal ganglia network. Frontiers in neural circuits, 7.

Lopez-Azcarate, J., Tainta, M., Rodriguez-Oroz, M. C., Valencia, M., Gonzalez, R., Guridi, J., Iriarte, J., Obeso, J. A., Artieda, J. and Alegre, M. (2010) Coupling between beta and high-frequency activity in the human subthalamic nucleus may be a pathophysiological mechanism in parkinson’s disease. J Neurosci, 30, 6667–77.

Mallet, N., Pogosyan, A., Marton, L. F., Bolam, J. P., Brown, P. and Magill, P. J. (2008a) Parkinsonian beta oscillations in the external globus pallidus and their relationship with subthalamic nucleus activity. J Neurosci, 28, 14245–58.

Mallet, N., Pogosyan, A., Sharott, A., Csicsvari, J., Bolam, J. P., Brown, P. and Magill, P. J. (2008b) Disrupted dopamine transmission and the emergence of exaggerated beta oscillations in subthalamic nucleus and cerebral cortex. J Neurosci, 28, 4795–806.

Marceglia, S., Fiorio, M., Foffani, G., Mrakic-Sposta, S., Tiriticco, M., Locatelli, M., Caputo, E., Tinazzi, M. and Priori, A. (2009) Modulation of beta oscillations in the subthalamic area during action observation in parkinson’s disease. Neuroscience, 161, 1027–36.

Maurice, N., Liberge, M., Jaouen, F., Ztaou, S., Hanini, M., Camon, J., Deisseroth, K., Amalric, M., Kerkerian-Le Goff, L. and Beurrier, C. (2015) Striatal cholinergic interneurons control motor behavior and basal ganglia function in experimental parkinsonism. Cell Rep, 13, 657–66.

McCarthy, M. M., Moore-Kochlacs, C., Gu, X., Boyden, E. S., Han, X. and Kopell, N. (2011) Striatal origin of the pathologic beta oscillations in parkinson’s disease. Proc Natl Acad Sci U S A, 108, 11620–5.

von Nicolai, C., Engler, G., Sharott, A., Engel, A. K., Moll, C. K. and Siegel, M. (2014) Corticostriatal coordination through coherent phase-amplitude coupling. Journal of Neuroscience, 34, 5938–5948.

Oswal, A., Beudel, M., Zrinzo, L., Limousin, P., Hariz, M., Foltynie, T., Litvak, V. and Brown, P. (2016) Deep brain stimulation modulates synchrony within spatially and spectrally distinct resting state networks in parkinson’s disease. Brain, 139, 1482–96.

Pisani, A., Bernardi, G., Ding, J. and Surmeier, D. J. (2007) Re-emergence of striatal cholinergic interneurons in movement disorders. Trends Neurosci, 30, 545–53.

Pittman-Polletta, B., Hsieh, W.-H., Kaur, S., Lo, M.-T. and Hu, K. (2014) Detecting phase-amplitude coupling with high frequency resolution using adaptive decompositions. Journal of neuroscience methods, 226, 15–32.

Priori, A., Foffani, G., Pesenti, A., Tamma, F., Bianchi, A. M., Pellegrini, M., Locatelli, M., Moxon, K. A. and Villani, R. M. (2004) Rhythm-specific pharmacological modulation of subthalamic activity in parkinson’s disease. Exp Neurol, 189, 369–79.

Rice, M. E., Patel, J. C. and Cragg, S. J. (2011) Dopamine release in the basal ganglia. Neuroscience, 198, 112–37.

Rubin, J. E. and Terman, D. (2004) High frequency stimulation of the subthalamic nucleus eliminates pathological thalamic rhythmicity in a computational model. Journal of computational neuroscience, 16, 211–235.

Sharott, A., Magill, P. J., Harnack, D., Kupsch, A., Meissner, W. and Brown, P. (2005) Dopamine depletion increases the power and coherence of beta-oscillations in the cerebral cortex and subthalamic nucleus of the awake rat. Eur J Neurosci, 21, 1413–22.

Shouno, O., Tachibana, Y., Nambu, A. and Doya, K. (2017) Computational model of recurrent subthalamo-pallidal circuit for generation of parkinsonian oscillations. Front Neuroanat, 11, 21.

Stein, E. and Bar-Gad, I. (2013) beta oscillations in the cortico-basal ganglia loop during parkinsonism. Exp Neurol, 245, 52–9.

Tachibana, Y., Iwamuro, H., Kita, H., Takada, M. and Nambu, A. (2011) Subthalamo-pallidal interactions underlying parkinsonian neuronal oscillations in the primate basal ganglia. European Journal of Neuroscience, 34, 1470–1484.

Terman, D., Rubin, J. E., Yew, A. and Wilson, C. (2002) Activity patterns in a model for the subthalamopallidal network of the basal ganglia. Journal of Neuroscience, 22, 2963–2976.

Threlfell, S., Lalic, T., Platt, N. J., Jennings, K. A., Deisseroth, K. and Cragg, S. J. (2012) Striatal dopamine release is triggered by synchronized activity in cholinergic interneurons. Neuron, 75, 58–64.

